# Robotic Stereotaxic System based on 3D skull reconstruction to improve surgical accuracy and speed

**DOI:** 10.1101/2020.01.04.894972

**Authors:** Phuong T. Ly, Alexandra Lucas, Sio Hang Pun, Anna Dondzillo, Chao Liu, Achim Klug, Tim C. Lei

**Affiliations:** The Department of Electrical Engineering, University of Colorado, Denver, CO 80204 USA; The Department of Physiology and Biophysics, University of Colorado, Anschutz Medical Campus, CO 80045 USA; State Key Laboratory of Analog and Mixed-Signal VLSI, Institute of Microelectronics, University of Macau, Macau, China

## Abstract

Some experimental approaches in neuroscience research require the precise placement of a recording electrode, pipette or other tool into a specific brain area that can be quite small and/or located deep beneath the surface. This process is typically aided with stereotaxic methods but remains challenging due to a lack of advanced technology to aid the experimenter. Currently, procedures require a significant amount of skill, have a high failure rate, and take up a significant amount of time.

We developed a next generation robotic stereotaxic platform for small rodents by combining a three-dimensional (3D) skull profiler sub-system and a full six degree-of-freedom (6DOF) robotic platform. The 3D skull profiler is based on structured illumination in which a series of horizontal and vertical line patterns are projected onto an animal skull. These patterns are captured by two two-dimensional (2D) CCD cameras which reconstruct an accurate 3D skull surface based on structured illumination and geometrical triangulation. Using the reconstructed 3D profile, the skull can repositioned using a 6DOF robotic platform to accurately align a surgical tool. The system was evaluated using mechanical measurement techniques, and the accuracy of the platform was demonstrated using brain phantoms. Additionally, small and deep brain nuclei were targeted in rodents for additional testing. The results indicate that this new stereotaxic system can improve the accuracy and speed of small-animal brain surgeries and reduce the failure rate of experiments.

## Introduction

Computer vision and robotics have made a significant impact on modern surgical practice. Several computer vision and robotic systems have already been used in hospital operating rooms to improve treatment outcomes and reduce the risks of surgical procedures ^1^. In order to increase dexterity and accuracy, some surgical robots were designed to perform motion tasks with multiple degrees of freedom for minimally invasive robotic surgeries (MIRS) ^2,3^, ranging from medium-sized laparoscopy and endoscopy to small-sized tissue grafting ^4,5^.

On the research side, stereotaxic (or stereotactic) surgery for small animals is an indispensable tool for many types of neuroscience studies ^6^. Stereotaxic surgeries are routinely performed in neuroscience laboratories for a variety of surgical procedures, including the creation of site-targeted lesions, injection of anatomical tracers, implantation of electrophysiology electrodes, and insertion of optical fibers or micro-dialysis probes ^7–9^. However, these stereotaxic surgical procedures are often time-consuming and prone to error due to the small size of the brain nuclei.

Current stereotaxic systems for small animals used in the laboratory are largely hand-driven and do not take advantage of modern electronic, mechanical and computer technologies ^10–12^. However, such advances have the potential to reduce surgical time and increase surgical accuracy. In addition, stereotaxic atlases used to estimate the 3D coordinates of the brain area with respect to anatomical landmarks on the skull are only available for very limited number of species and age groups ^11^. Mechanical-based injections or surgical procedures have largely not been automated, and are prone to error, with success rates as low as 30% when small and deep brain areas need to be precisely targeted.

In recent years, there have been efforts to develop improved stereotaxic systems that automate stereotaxic surgeries to reduce human error, improve surgical accuracy and save time. Pak et al., developed an automated craniotomy system which controls a motorized drill based on impedance measurements to open a cranial window as small as several millimeters with high repeatability ^13^. Although this system is robotically and electronically controlled, it is not a full stereotaxic system, lacking the capabilities to perform stereotaxic positioning and automatic insertion of surgical tools. The system was only designed to retrofit to existing manual stereotaxic systems and to open holes on the skull automatically with high spatial precision. Neurostar, on the other hand, developed a robotic stereotaxic system in which the three translational actuators were motorized, and an electronic brain atlas was added to the control computer to guide the experimenter during stereotaxic surgeries ^14^. This system can also be optionally equipped with an electronic drill and microinjector to provide automatic skull drilling and dye injections. However, the Neurostar system only provides three motorized translational actuators, and rotational positioning remains to be manually controlled. This makes it impossible for the system to automatically align the animal to the “skull-flat” position, meaning that the skull profile is parallel to the laboratory coordinate, and is critical for precise nucleus targeting to avoid blood vessels or other important brain regions. The system also uses a pair of 2D CCD cameras to identify the single Bregma landmark on the skull and guide the surgical tool, but the design lacks the needed hardware, such as structured illumination light sources, for a full 3D skull profile reconstruction. Another robotic stereotaxic surgical system was developed by Brainsight ^15^. The Vet robot of Brainsight uses a full 6 degree-of-freedom (6DOF) robotic arm to guide a surgical tool to a brain region. The system is also integrated with MRI or CT images for automatic stereotaxic guidance. The Vet robot is equipped with two CCD cameras for 3D positioning registration and a 200 µm positioning accuracy was demonstrated on their system. Comparing to the Neurostar system, the Brainsight system’s full 6 degree-of-freedom robotic arm allows surgical insertions at all angles. However, the stereo camera system still lacks the capability to reconstruct the skull profile in 3D and only a single target point can be identified. Another disadvantage of using a full 6 degree-of-freedom robotic arm is the high cost of ownership, which makes the system difficult to adapt for many neuroscience laboratories.

The objective of this study was to develop a fully automated stereotaxic system for small animals. The system will significantly improve the surgical speed and success rate of brain surgeries due to the improved spatial resolution and accuracy for both measurement and overall manipulation. The system is equipped with a 3D skull profiler which uses structured illumination and geometrical triangulation to map the skull surface of a small rodent with high spatial (sub-millimeter) precision. In addition, a full 6DOF robotic platform will provide a large translational and rotational range of motions to position the animal for precise stereotaxic procedures. In this paper, the design and construction of the system is discussed in detail and the accuracy evaluation of the system using both mechanical measurement and actual stereotaxic injection on brain phantoms and anesthetized rodents are also demonstrated.

## Results

The main goal of this project was to design and build a stereotaxic device that can perform surgery on small animals such as rodents automatically and without human intervention, thus leading to increased targeting accuracy especially for small and deep brain nuclei. More specifically, we intended to address two areas of inaccuracy that many existing devices have: 1) The acquisition of measurements needed to obtain skull-flat and 2) inaccuracies with the movement of the various axes to adjust, rotate, and tilt the animal’s skull into the correct position.

To address the first area of concern (measurements of skull-flat), we developed a computer vision system that performs 3D scans of an animal’s skull ^16,17^. The computer vision system is a 3D skull profiler that can scan the rodent’s skull using a video projector to project line patterns which are then imaged by two regular CCD cameras carefully attached on either side of the projector ^18^. The acquired images can be used to calculate a 3D skull profile with sub-millimeter spatial resolution based on the techniques of structured illumination and geometrical triangulation ^19^.

To address the second area of concern (precision of movement), we developed a stereotaxic platform that is based on a hexapod (Stewart) design ^20–23^. A Stewart platform consists of two plates that are connected to each other via six motorized axes. Lengthening or shorting these six axes in a synergistic way will allow the top plate to rotate, tilt, or move against the stationary bottom plate with six DOF (X, Y, Z, roll, pitch, yaw). Based on the spatial information reconstructed from the 3D skull profile, the animal skull can be moved to a desired location according to the specific brain area calculations derived from brain atlas coordinates.

Figure 1 shows both the schematic diagram and the actual photographic image of the automated robotic stereotaxic system illustrating the major components required for the system.

**Figure 1.**
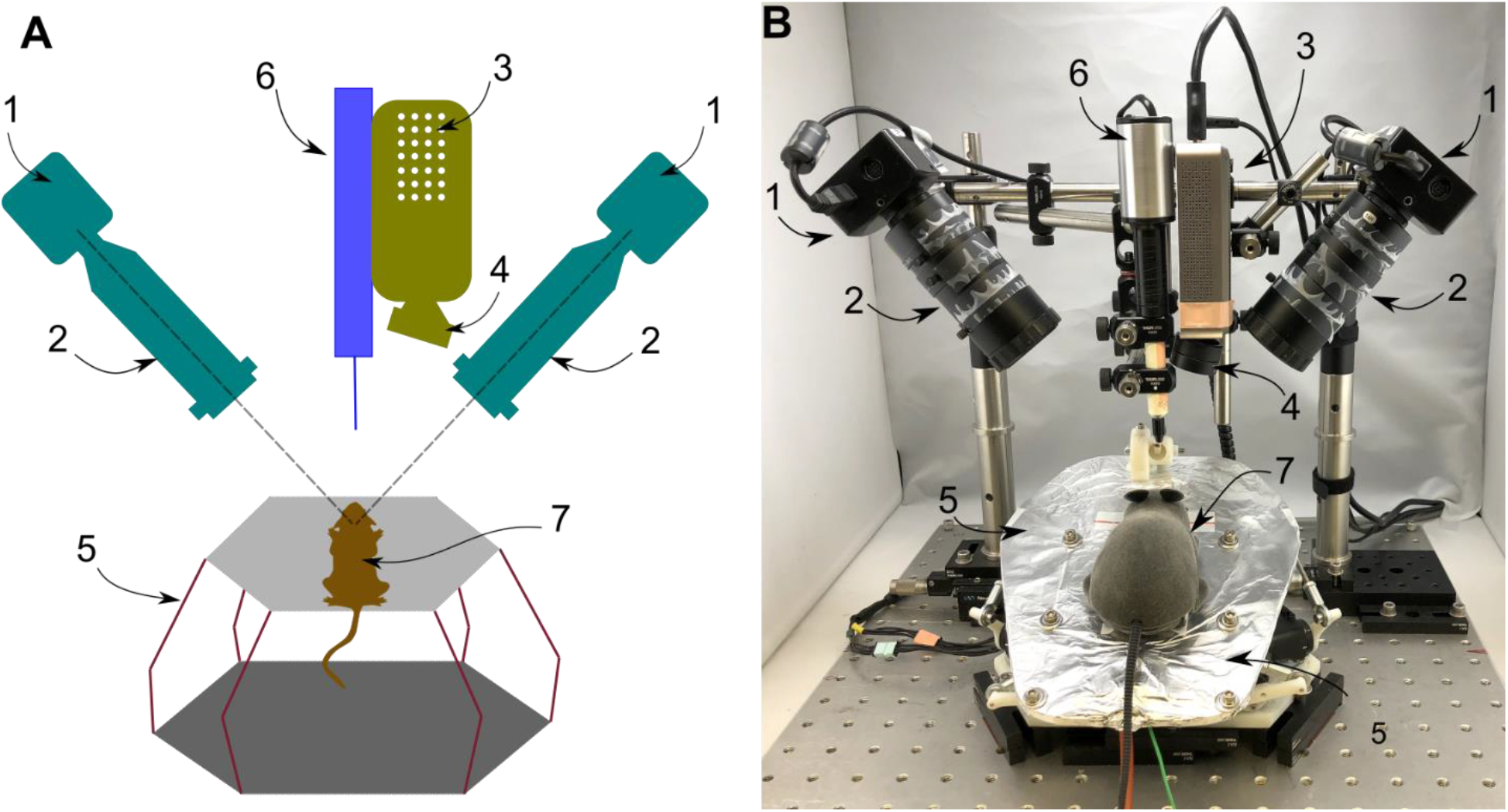
Fully automated robotic stereotaxic system based on 3D structured illumination and geometrical triangulation. **A.** Schematic diagram of the stereotaxic system illustrating the crucial components of the system. Two cameras (1) mounted onto the two sides of the center axis were equipped with zoom lenses to focus on the rodent’s skull (7). A video projector (3) was used to project structured light patterns onto the skull surface. A 75 mm bi-convex lens with a small slanted angle was used to focus the projection plane of the structured image onto the skull. A 6 degree-of-freedom (3 translational and 3 rotational) robotic platform (5) was used to secure the rodent and allowed positioning the animal’s head with sub-millimeter spatial resolution. A surgical device (6) such as a nanoliter injector can be placed along the center axis to perform stereotaxic surgical procedures. **B.** Photographic image of the actual automated stereotaxic system.

### 3D skull profile reconstruction using structured illumination

The skull of a rodent can be reconstructed in 3D space through the technique of structured illumination. Figure 2A shows example images of the skull illuminated with vertical and horizontal line patterns taken by the left and right 2D CCD cameras. A total of 42 photos were taken by each camera with the spatial frequencies varying from 0.025 to 25.6 lines/millimeter for 3D reconstruction. According to the rule of optical projection, vertical displacement on the rodent’s skull in turn creates lateral displacements in the acquired 2D images. These lateral displacements can be used to estimate the vertical displacement of each point on the skull surface using geometrical triangulation. To correctly calibrate the camera systems and obtain sub-millimeter spatial reconstruction, 3D printed calibration targets and optical calibration targets with known dimensions were first imaged to estimate the correct parameters for the focal lengths of the CCD cameras and the relative positional parameters for the cameras and the projectors. Through this calibration process, it was determined that the 3D reconstructed skull profile can achieve a spatial resolution of 98.5±4.5 µm.

**Figure 2:**
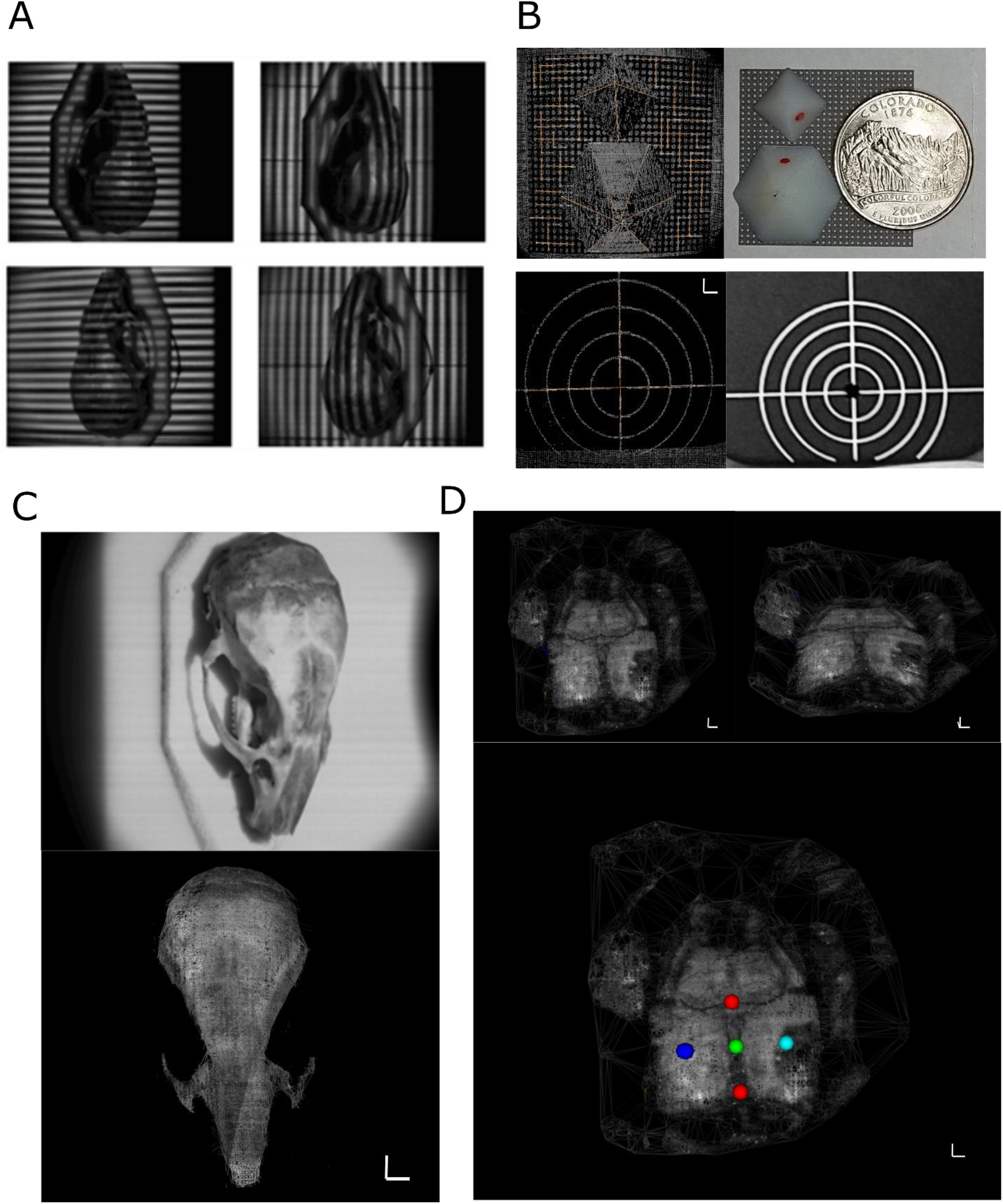
3D skull profile reconstruction using structured illumination. **A.** A Mongolian gerbil skull was illuminated by example vertical and horizontal lines observed by the left (top) and the right (bottom) cameras. **B.** Reconstructed calibration standards using structured illumination (top left) and the 3D printed calibration standard in pyramided shapes (top right). Reconstructed optical target (bottom left) and the calibrated optical target (bottom right) for calibrating the reconstruction routine to obtain sub-millimeter spatial resolution. **C.** A regular 2D image taken by the left black and white CCD camera (top) and the reconstructed 3D skull profile (bottom) of a Mongolian gerbil skull. The reconstructed 3D points of the skull profile were colored with greyscale intensity to visualize stereotaxic landmarks (Bregma and Lambda). **D.** Reconstructed 3D skull profile of an anesthetized Mongolian gerbil in two different view angles clearly showing the exposed skull and skin (top). The Bregma (top red dot) and Lambda (bottom red dot) was used to identify and to estimate the center intersection point (green dot) and two side points (left blue and right cyan dots) 4 millimeters perpendicular to the center connecting line (bottom). Note: all scalebars are 1 mm in length.

The 3D profiler was used to reconstruct the profile of an *in-vitro* Mongolian gerbil skull. The 2D normal CCD image and the reconstructed 3D profile of the gerbil skull are shown in Fig. 2C. The reconstructed 3D skull profile was superimposed with greyscale intensity obtained from a normal 2D skull images for better visualization. The stereotaxic landmarks and bone sutures can be clearly identified from the reconstructed 3D skull profile. The reconstructed skull profile has very few missing 3D points on the skull surface. The structures on either side of the skull surface were not reconstructed, and this is because only one camera can see a given side of the skull, making geometrical reconstruction not possible for these points. However, the inability to reconstruct the side surfaces was not important for our purposes since the normal plane for the rodent’s skull can be accurately determined using the top skull surface alone.

After confirming the functions and accuracies of the computer vision 3D profiler with an *in-vitro* skull, an anesthetized Mongolian gerbil was then used for additional system evaluation. The gerbil was secured to the robotic platform using dental cement and a headpost attached to the top of the robotic platform (Figure 3H). The scalp of the animal was surgically opened, exposing the top surface of the skull including Bregma and Lambda, two skull suture points which are commonly used in stereotaxic surgeries (red dots in figure 2D). The surface of the skull was scanned by the 3D profiler without the use of image enhancer, fiducial markings or other treatment methods. The reconstructed 3D surface profile is shown in Fig. 2D in two different viewing angles (top figure). The skull surface was well-reconstructed showing details including the stereotaxic landmarks and part of the exposed scalp. Using our custom control software, the reconstructed 3D skull profile can be manipulated on the computer screen in real-time using a computer mouse, and the Bregma and Lambda points can be confirmed by the user. Based on these two stereotaxic landmark points, the mid-point between them, plus two additional points 4 mm lateral to the midline were estimated automatically by the software. Based on these points, two perpendicular 3D lines are determined defining a normal surface plane for the gerbil skull. This normal plane can then be used to move the robotic platform to the “skull-flat” position, in which Bregma and Lambda are at the same height with an error of less than 100 µm, for subsequent alignment and stereotaxic procedures.

**Figure 3:**
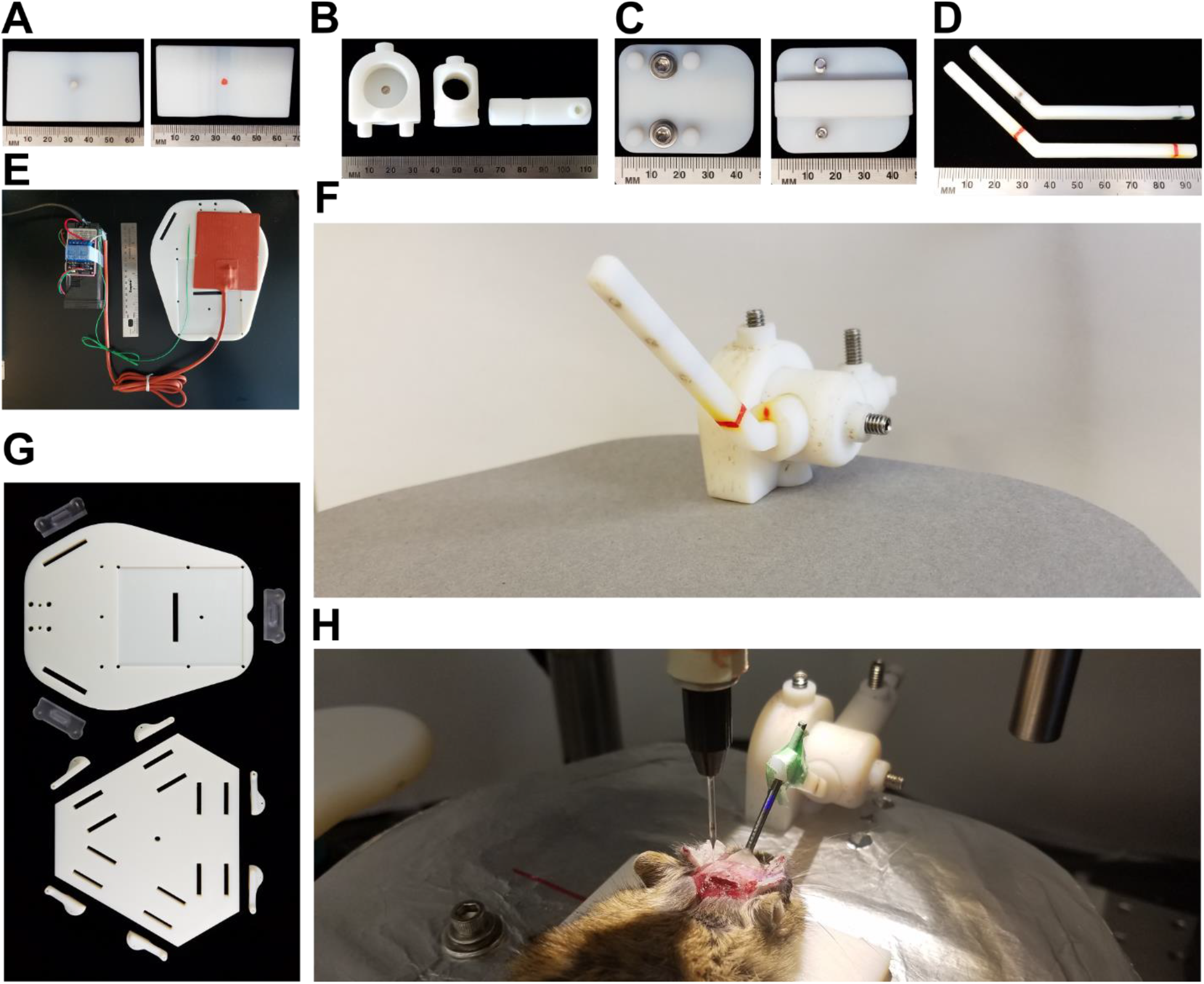
Components and add-ons of the six degree-of-freedom robotic platform. **A.** A 3D printed head rest that can be optionally attached to the platform to better position the animal’s head for stereotaxic surgery. **B to D.** 3D printed sub-components of a rotatable and extendable manipulator (**F**) to hold a metal bar that can be fixed to the skull for securing the animal. **E.** A temperature heating pad controlled by a PID controller can be embedded into the top plate of the robotic platform to maintain the body temperature of the rodent. **G**. 3D printed top and bottom plates with attaching parts and short arms for motors of the robotic platform in which the top plate has several mounted holes to allow add-ons to be installed.

### Full six degree-of-freedom robotic stereotaxic platform developed using 3D prototyping technology

The full six degree-of-freedom robotic stereotaxic platform was manufactured using rapid prototyping technology. The majority of the components were created using 3D printing technology and were designed using Solidworks 3D CAD design software (Dassault Systèmes, Vélizy-Villacoublay, France). The components were then printed using a high-resolution 3D printing system (Objet30, Stratasys, Rehovot, Israel) with a spatial resolution of 28 µm. Rapid prototyping manufacturing technology allows the platform to be easily configured to hold different animal species and sizes. Additional accessories are also required for stereotaxic work to succeed with live animals; for example, many animal species and/or surgical procedures require that the animal maintain physiological temperature. Also, the animal’s head needs to be immobilized so it remains stable and in position during the surgeon’s handling and manipulation. Depending on the brain area and investigator’s experimental requirements, this fixation may occur via ear bars, a bite bar, a head post, a head rest, or a combination of these methods. Figure 3 shows some custom add-ons that were manufactured with 3D printing prototyping technology. Figure 3A is a custom head rest that is designed for medium sized rodents such as Mongolian gerbils. In addition, many stereotaxic devices use ear bars to secure the rodent on the platform; however, these cannot be used in some experiments, for example those involving the auditory pathway. For that reason, a rotatable and extendable manipulator was designed to hold a head post which can be secured with dental cement to the skull of the rodent. Figures 3B - D show the sub-components of the manipulator which can be assembled to attach to the top plate of the robotic platform to secure the animal, as shown in Figure 3H. Temperature control is essential to the survivability of the rodent and the rapid prototyping technology allowed for integration of a heating pad with temperature control to the platform. Figure 3E shows the heating pad controlled by a proportional-integral-derivative (PID) controller with a thermostat placed underneath the animal to ensure that a constant temperature of 37°C is maintained throughout the entire stereotaxic surgery. The heating pad can be embedded into the top plate of the platform seamlessly to provide maximum compatibility. Figure 3G shows the top and bottom plates of the robotic stereotaxic platform with constructive parts also prototyped using 3D printing technology. The technology allows custom mounting holes and placement grooves for the top plate to easily mount a variety of components for securing the animal or other life-support related installments.

The robotic platform was first characterized using mechanical calibration techniques. Considering the radius of a small rodent’s skull, which is in the order of 20 mm, and the typical dimensions of target brain regions inside the rodent’s brain which can be as small as 0.2 mm, translational and rotational accuracies of 200 µm and 0.5 degrees, respectively are required to accurately target any given brain region. In addition, the translational and rotational motions of the robotic platform must be linear to ensure smooth positioning for stereotaxic surgeries. Three mechanical dial gauges (Model 25-611, L. S. Starrett, Athol, MA, USA) and a digital 9 degree-of-freedom inertial measurement unit (IMU) chip (MPU9150A, InvenSense, San Jose, CA, USA) were used to measure both the translational displacements and the rotational angles of the robotic platform against the desired translational distances or rotational angles issued by the control computer. The three translational gauges were mounted against the platform perpendicular to one another for all three axes, and the IMU chip was directly mounted at the center of the top plate to measure the rotational angles. Figure 4 shows some selected calibration curves (the z linear axis and the pitch angle) to demonstrate the linearity in both translational and rotational motions. Fig. 4C compares the desired and actual movements in all three translational and rotational axes. The overall error is less than 4.4 %. Using calibration tools, the robotic platform was determined to have a full translational distance of ± 15 mm with an accuracy of ± 0.25 mm, and a full rotational angle of ± 20° with an angular accuracy of better than ±0.1°, as summarized in Table 1.

**Table 1:** Maximum translational and rotational ranges and resolutions of 6 DOF robotic platform

| Top platform | Measured ranges and accuracies |  |
| --- | --- | --- |
|  | *Translational* | *Rotational* |
| Maximum range | $\pm 15$ mm | $\pm 20^\circ$ |
| Resolution | $\pm 0.25$ mm | $\pm 0.1^\circ$ |

**Figure 4:**
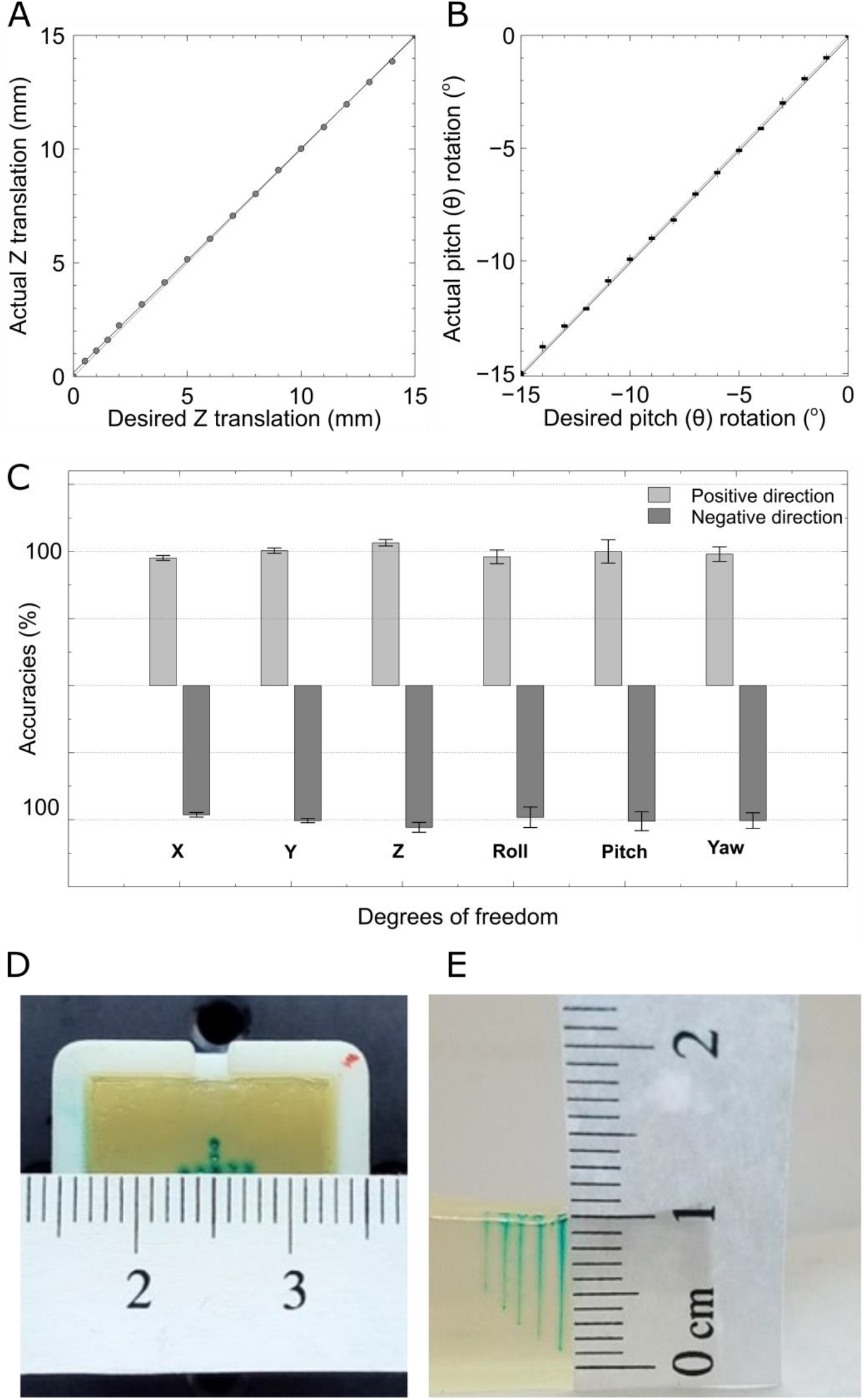
Positioning accuracy of the robotic stereotaxic platform **A.** Desired Z translational positions against the actual Z translational positions by the platform. **B.** Desired pitch rotational angles against the actual pitch rotational angles by the platform. **C.** The estimated accuracies in all six degree-of-freedom motions for the robotic platform. **D.** A cross pattern with an inter-spacing of 1 mm was injected into an agar brain phantom using the stereotaxic platform. **E.** Longitudinal injections, performed with the stereotaxic platform, with a depth difference of 1 mm were quantified in the agar brain.

The robotic platform was also evaluated using a brain phantom made from agar to demonstrate its positioning accuracy. The robotic platform was first programmed to print a “cross” pattern with an inter-spacing of 1 mm into the brain phantom, as shown in Fig. 4D. The diameter of the glass micropipette tip used in the injection was estimated to be ~100 µm and there is no observable derivation of the printed pattern on the brain phantom. Injection depth accuracy was then demonstrated via vertical injections with a 1 mm depth difference as shown in Fig. 4E, and the injection printing again has no observed derivation.

### Stereotaxic in-vivo injections into a deep brain nucleus

To test the system’s performance in actual stereotaxic procedures, an anesthetized gerbil was placed on the top plate of the robotic stereotaxic platform and secured via a head post attached to a customized manipulator on the top plate, as shown in Fig. 5A. The top scalp of the gerbil was incised exposing the skull, and structured illumination scanning was used to construct the 3D skull profile on the computer. Bregma and Lambda landmarks were identified and selected on the computer to determine the skull “normal” surface. The “skull-flat” position was achieved by calculating the required translational and rotational displacements of the skull normal surface and translating into a computer command to move the robotic platform with the gerbil to the determined position. From skull-flat, the animal together with the top plate was tilted via an additional rotational displacement of −20 degrees pitch to facilitate access to the desired brain area, and additional translational displacements of 4 mm posterior, then 0.8 mm and 0.6 mm laterally to the left or right of Lambda were initiated. These displacements were estimated based on a gerbil brain atlas (Radke-Schuller et al, 2016) to target the MNTB in the brainstem. After drill positions were marked and two small holes were drilled manually, the skull was re-scanned by the 3D computer vision skull profiler. The robotic platform then was commanded to compensate for any positional errors induced by the drilling process. A glass pipette with tip diameter ~100 µm, which was filled with either one of two fluorescent dyes: cascade-blue (ThermoFisher, Waltham, MA; for the left hemisphere injection) or micro-ruby (ThermoFisher, Waltham, MA; for the other right side), was inserted to the gerbil head through each one of the holes and advanced to a at a depth of 7.5 mm by gradually raising top plate of the robotic platform through a series of commands towards the animal, as shown in Fig. 5B. Note that this procedure is unusual and was used for demonstration purpose with our prototyping setup, and the main purpose of these dye injections was to test the accuracy of the platform and therefore we aimed to avoid additional third-party technology that might obscure our measurements. During typical laboratory use of the platform, the investigator would most likely use an additional motorized axis for lowering the electrode for better controller and higher accuracy. After the injections, the animal was sacrificed, and the brain was extracted and sectioned for fluorescence imaging. Fig. 5C shows a coronal section with a yellow and a blue tract, showing the path of the electrode to the target area the medial nucleus of the trapezoid body (MNTB). The insert in figure 5c show this area marked in a brain atlas sketch. This simulated neuroscience animal stereotaxic surgery confirms that the system can inject to the intended brain region precisely with very minimal user inputs.

**Figure 5:**
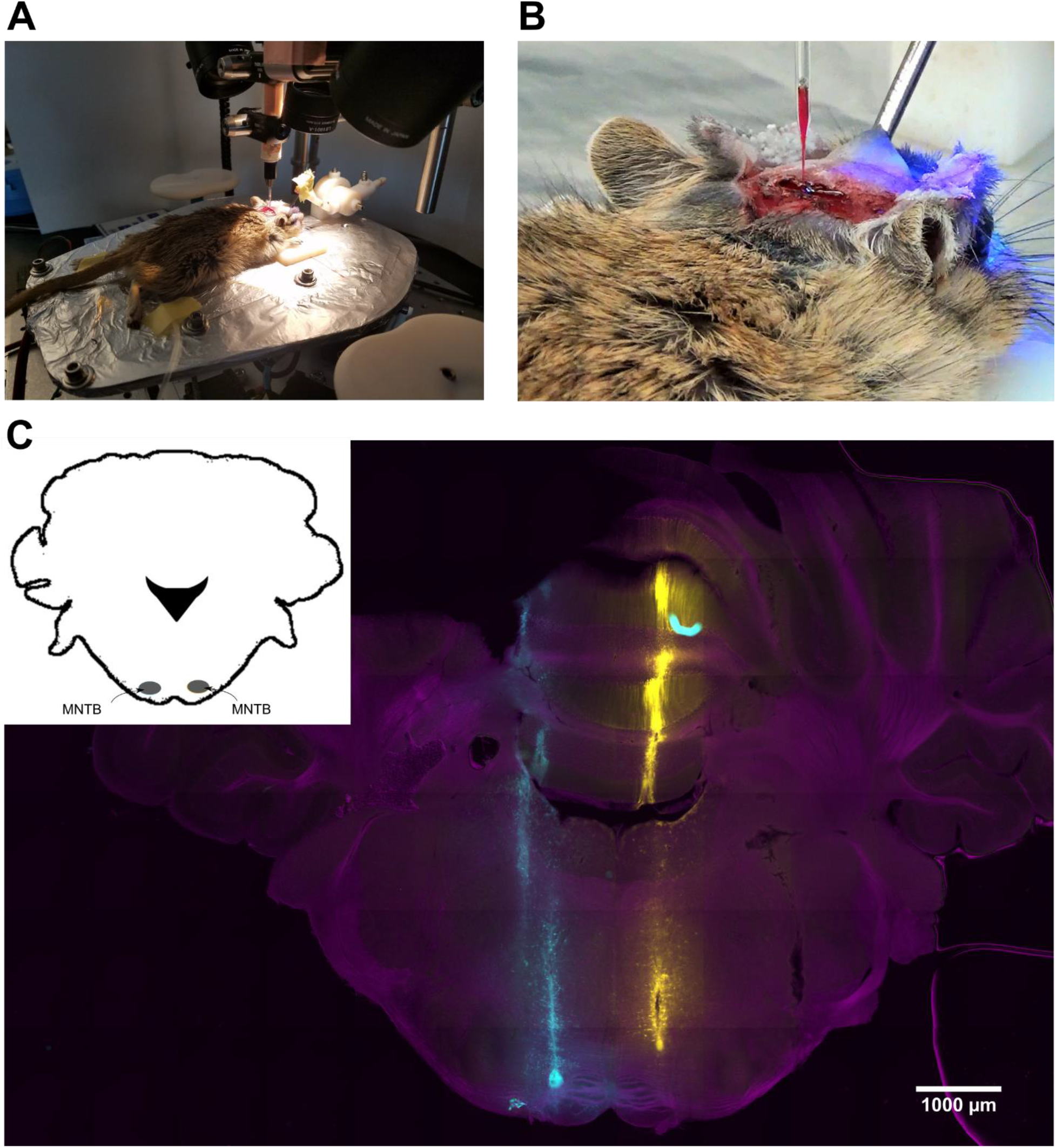
Fully automatic stereotaxic brain surgery on a Mongolian gerbil. **A.** The exposed skull of an anesthetized Mongolian gerbil was first 3D scanned and automatically positioned for stereotaxic injection to the ventral end of the left medial nucleus of the trapezoid body (MNTB) (blue tracer) and the dorsal end of the right MNTB (yellow tracer), deep within the brain. **B.** Close-up view of the stereotaxic injection with the glass micropipette inserted into the animal’s brain. **C.** Fluorescence imaging of an excised brain slice showing the successful targeting of the MNTB region. Upper left is the schematic brain atlas showing the locations of the two MNTBs.

A video showing the entire automatic stereotaxic alignment process including image taking, 3D skull profile reconstruction, picking the skull landmarks and computer-guided platform moving to achieve skull-flat position is provided in the supplementary information. Note that the video was 5x time-accelerated to save the reader’s time and the length of the video is 63 seconds, making the entire process to be less than 5 and a half minutes.

## Discussion

In this paper, we demonstrated that an automatic robotic stereotaxic system was realized by combining a 3D computer vision optical skull profiler based on the techniques of structured illumination and geometrical triangulation, and a 6DOF robotic platform based on the Stewart design. Through this combination, the skull profile of a small rodent can be reconstructed in computer space with a high degree of accuracy. The 6DOF robotic platform can then be instructed by the positioning estimations to move the rodent’s skull to the correct position for stereotaxic surgeries. To the best of our knowledge, this is the first time a 3D optical profile is being used to capture the 3D point cloud of a rodent’s skull for stereotaxic purposes. While other existing robotic stereotaxic systems use single or multiple CCD cameras, these cameras are mostly used to determine the coordinate of a single or multiple fiducial spheres or landmarks. The advantage of using our approach is that no preparation or device mounting is required to provide a full 3D reconstruction of the skull profile for precise stereotaxic positional identification. Our Stewart based platform design provides full 6DOF motions in all translational and rotational axes with a good range of movements (±15 mm and ±20°) and high precision (±0.25 mm and ±0.1°), which is sufficient to cover all translational and rotational needs for small rodent stereotaxic procedures. In addition, the range of platform movements is related to the variation lengths of the six arms, thus the platform can be designed in terms of platform size and arm lengths to accommodate other animal dimensions.

Another design advantage is that the system is constructed with relatively low-cost components, making it highly affordable and suitable for wide use. The 3D optical skull profiler is constructed using two low-cost CCD cameras with two 10x tele lenses and a commercial computer projector with a low-cost 75 mm biconvex lens. This design does not require special time-of-flight 3D cameras or other special optical components for the 3D reconstruction ^24,25^. The robotic platform was built using 6 low-cost digital servos and the other components were 3D printed using rapid prototyping technologies. Originally, six linear translational actuators were used to realize the Stewart platform design, but these required good spatial resolution which drastically increased the building costs of the platform. In contrast, digital rotational servos are more economical, and can be used to achieve precise movement outcomes through trigonometric relation as illustrated in the supplementary information. Therefore, the digital servos were chosen to build the rotational platform, allowing the device to remain accessible while maintaining high spatial positioning accuracy. Additionally, our system is designed to move the animal together with the robotic platform while other robotic stereotaxic systems keep the animal stationary. This allows the robotic platform to achieve accurate stereotaxic results but with a significant cost reduction.

An increasing number of experimental approaches in neuroscience require the precise placement of a recording electrode, injection pipette or some other tools into a specific brain area that can be quite small and/or located deep beneath the surface. Reaching these brain areas with traditional methods and devices can be challenging for several reasons. First, some brain nuclei are less than 0.5 mm in diameter and may be as deep as 7-10 millimeters in typical rodent species, requiring a target accuracy of better than 1 degree. Second, investigators try to minimize the size of the opening holes in an animal’s skull through which a tool is advanced to minimize surgical trauma from the intervention. Third, jaws, face, or ears additionally limit the locations on the head where craniotomies can be performed, such these practical reasons, most craniotomies use a dorsal approach. This lack of sophisticated technology is especially unacceptable for many other types of experiments that target small and deep brain areas (midbrain, brain stem, thalamus, subregions of hippocampus, and many other types of non-surface structures). As a result, many in-vivo manipulations either require a significant amount of experience by the experimenter, and/or have a significant failure rate. Failed experiments are costly in terms of wasted investigator time, research animals and materials. However, even in cases where a lab has the expertise to target the desired brain area with a relatively high success rate (for example, because they employ a very skilled student/postdoc/lab tech), this is problematic since the success depends on that person and their “magic touch” – effectively reducing the reproducibility of these experiments for all other labs that don’t have access to that person. Even the same lab may have trouble reproducing their own experiments once that person with the “magic touch” leaves. The idea behind the device described here is to eliminate qualitative aspects of stereotaxic procedures, such as the skill and experience of the operator as much as possible, and to replace these qualities with precisely measured and precisely repeatable automated procedures. Neuronavigational features can be added to the device in the future, which would increase the automatism and the repeatability of stereotaxic work with this device further.

The system can be further enhanced in the following areas. The current software was written with the Python programming language, and its mathematic calculations can be slowed by the interpretative nature of the programming language. In addition, single core computing was only used in the 3D reconstruction algorithm and improvements in calculation speed can be achieved using multiple CPU cores or GPU acceleration to reduce the time needed for the reconstruction ^26^. The current software requires users to visually identify the Bregma and Lambda points on the 3D reconstructed skull profile and this identification process can further be automated using image processing techniques in the future. Holes for injections were still manually drilled, and additional drilling add-ons can be installed to allow automatic drilling. Installing an impedance-based sensor on the drill attachment and platform like the design of Pak et al. will allow precise drilling of the skull without damaging brain tissue ^13^.

## Conclusion

A new type of stereotaxic system for small animal brain surgeries has been developed by combining a 3D computer vision sub-system and a 6DOF robotic platform. High resolution 3D reconstruction of an animal skull has been demonstrated. No special 3D camera or hardware was required, but a series of images were captured by two regular computer cameras mounted on both sides of the skull. A series of structured patterns were projected onto the animal skull using a video projector to increase optical contrast and allow high resolution 3D capture on a relatively featureless skull structure. Six low-cost digital servo motors were used and controlled the 6 extendable arms to allow placement of the top platform in all 3 translational and 3 rotational positions with few limitations. Both the 3D camera sub-system and the robotic platform were characterized to have sub-millimeter and sub-degree spatial resolution suitable to precisely target a small neural region within the animal’s head. A simulated stereotaxic surgery using an anesthetized Mongolian gerbil has confirmed that the MNTB can be accurately targeted using the system. We anticipate that this new development will help the advancement of neuroscience research through an increased success rate for stereotaxic surgeries and reduced surgical time to increase animal survivability.

## Methods and System Design

### Optical structured illumination for precision 3D skull mapping

A video LED projector (S1, ASUS, Taipei, Taiwan) was mounted slightly offset from the middle longitudinal axis of the entire setup because commercial projectors are designed to project videos in a tilted upward position. Since the video projector is designed to project videos meters away, a 75 mm bi-convex lens (LB1901-A, Thorlabs, Newton, NJ, USA) was placed in front of the projection lens to focus the projecting images at a much shorter distance onto the rodent’s skull. Two 2D CCD cameras (BCE-B013-U, Mightex, Pleasanton, CA, USA) were attached on both sides of the projector to capture skull images illuminated with projected line patterns. Two 10x zoom lenses (MLH Macro 10X, Computar, Las Vegas, NV, USA) were also mounted onto the two CCD cameras to reduce field of view onto the rodent’s skull and maximize the image resolution for the 3D reconstruction. The two CCD cameras have a pixel density of 1280 × 1024 covering a field-of-view sized approximately 50 × 40 mm which translates to a lateral spatial resolution of 30 × 35 mm for the 3D reconstructed profile.

A series of horizontal and vertical black-and-white lines with increasing spatial frequencies (structured illumination) were projected onto the rodent’s skull, and the two CCD cameras were used to capture 2D images of the skull covered with projected lines from both sides. Using these captured 2D images with structure-illuminated patterns, a unique binary spatial code can be constructed to identify all points on the skull surface. Based on the spatial code, the rodent’s skull surface can be reconstructed in 3D using geometrical triangulation in which the projected lines are laterally displaced when projected onto a non-flat surface and the degree of lateral displacement is linearly proportional to the vertical corrugation of the rodent’s skull. Therefore, a surface point *P* in the 3D space can be estimated by

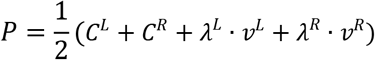

where *ν*^*L*^ and *ν*^*R*^ are the viewing vector pointing towards the point *P* for the left and right cameras; *C*^*L*^ and *C*^*R*^ are the 3D coordinates of the center points of the left and right cameras; and *λ*^*L*^ and *λ*^*R*^ are parameters can be calculated based on geometric triangulation. Detail discussion for the equations to calculate *λ*^*L*^ and *λ*^*R*^, and the 3D structure illumination methods can be found in the supplementary information. A custom Python program was written to process the captured 2D images and reconstruct the 3D skull profile. A total of 42 vertical and horizontal line patterns were projected onto the rodent’s skull and captured by each of the CCD cameras with a frame rate of 25 fps. All the images were captured in standard laboratory lighting conditions with no light shielding employed. The exposure time for each image was 20 ms with an average of 8 frames to improve image contrast and to remove optical noise induced by the projector and room lighting. The overall image capture time was approximately 86 seconds. An Intel Core i5-4430 CPU @ 3.00GHz with 16.0 GB RAM commercial desktop computer was used for the reconstruction process along with a custom Python program. The processing time to reconstruct the 3D profile was approximately 9 seconds.

### Full 6 degree-of-freedom robotic platform

The robotic platform provides a full 6 degree-of-freedom - 3 translational and 3 rotational - movements to allow precise positioning of the rodent’s skull. The platform is based on the Stewart design in which six motorized arms are attached to a moveable top plate and a stationary bottom plate. The movements of the 6 arms are coordinated to give the top plate a full 6 degrees of motions. This is different from other conventional Stewart designs in which 6 linear actuators are used. The robotic platform used in this project was built with 6 digital rotational servo motors (AX-12A, Robotis, South Korea), and each was connected to a pair of short and long arms which formed a semi-triangle with a rotational pivot. The short and long arms act as the two known edges of a semi-triangle and through rotating the servo angle, the hypotenuse edge of the semi-triangle can be shortened or lengthened to provide the desired linear extension for coordinated platform motion. Details of the control algorithms for the platform can be found in the supplementary information. This design allows the platform to have precise sub-millimeter translational and sub-degree rotational positioning accuracies and at the same time reduces the cost of ownership using low cost rotational servo motors. In the current system, the top and bottom plates were initially designed with Solidworks mechanical design software, then were 3D printed by a 3D plastic printer (Objet30, Stratasys, Rehovot, USA) with a printing spatial resolution of 28 µm. The top plate was also embedded with a heating pad which was controlled by a PID controller to maintain 37°C body temperature for the rodent. Multiple accessories, such as ear and bite bars, can also be mounted on the top plate for securing the animal on the platform.

### Fully automatic stereotaxic skull alignment and positioning

Fully automatic stereotaxic surgery can be achieved by coordinating the 3D reconstructed skull profile obtained from the computer vision system with the movements of the 6 DOF stereotaxic platform through the custom Python control program. A small rodent can be securely supported on the top plate of the robotic platform either using a pair of ear bars or a metal head post fixed to the rodent’s skull, and the temperature of the rodent can be maintained through a PID-controlled heating pad. After the head is shaved and the skull is exposed, a series of structured illuminated images taken by the two CCD cameras will create the 3D skull profile which can then be reconstructed by the software routine. Users are prompted to select the Bregma and Lambda landmark positions by clicking on the 3D reconstructed skull profile which is rendered in 3D viewing window on computer screen. Based on the 3D coordinates of Bregma and Lambda, the mid-point between these landmark points can be automatically estimated. In addition, two points that are 4 millimeters away from this mid-point and are perpendicular to the line connecting Bregma and Lambda points will be estimated. Thus, these five points (Lambda, Bregma, mid-point, two side points) form two perpendicular lines to define a 3D surface normal plane for the rodent’s skull in space. Using this surface normal plane, the robotic plane can be translated and rotated to achieve the “skull-flat” position, where the Bregma and Lambda landmarks are at the same height level in the robotic plane. At this point, the users can enter the desired translational and rotational displacements referenced to the landmarks of the skull for the robotic platform moving to any desired location ^11^.

### Animal protocol and procedures for stereotaxic evaluation

All experimental procedures complied with all applicable laws and NIH guidelines and were approved by the University of Colorado IACUC. All experiments were conducted in adult Mongolian gerbils (Meriones unguiculatus). Animals were first anesthetized with a mixture of ketamine-xylazine (60mg/kg-5mg/kg), and a maintenance dose (25mg/kg-5mg/kg) was given following complete anesthesia to maintain the anesthetized state. Once the animal was properly anesthetized, the fur over the skull was shaved off and the underlying skin sanitized with a disinfectant. Skin and muscle overlying the skull were removed and a craniotomy was made in the skull at 0.8 mm and 0.6 mm lateral and 4 mm posterior of lambda using a dental drill. The coordinates used here allowed us access the auditory brainstem for dye injections into the MNTB. Dye injections were made using a Nanoliter injector (World Precisions Instruments, Sarasota, FL). 32.2nL of dye was injected at a depth of 7.5mm once every 30 seconds and 8 injections were made at each location. After all injections were concluded, the animal was given an overdose of pentobarbital (0.03 mL/g) and perfused transcardially with phosphate buffer solution (PBS) and 4% Paraformaldehyde (PFA). Once perfusion was complete, the brain was fixed in 4% PFA overnight, then removed and placed into 4% agar. The brainstem and cerebellum were sliced coronally in 100 µm sections using a vibratome (Leica VT 1000s, Nussloch, Germany). The sections were then stained with a 1:100 concentration of Neurotrace Nissl stain (ThermoFisher, Waltham MA, 640/660 deep red fluorescent Nissl stain) diluted in antibody solution (ABS) ^27^. The slices were mounted on slides with Fluoromount-G (Diagnostic BioSystems, Pleasanton CA) and imaged with an Olympus FV1000 (Tokyo, Japan) confocal microscope using the laser lines of 405nm 555nm, and 647nm to image the Nissl (cell body indicator).

## Data availability

The data that support the findings of this study are available from the corresponding author upon reasonable request.

## Reference

1. Lanfranco, A. R., Castellanos, A. E., Desai, J. P. & Meyers, W. C. Robotic Surgery: A Current Perspective. Ann. Surg. 239, 14–21 (2004).

2. Hagn, U. et al. DLR MiroSurge: A versatile system for research in endoscopic telesurgery. Int. J. Comput. Assist. Radiol. Surg. 5, 183–93 (2010).

3. Hannaford, B. et al. Raven-II: An open platform for surgical robotics research. IEEE Trans. Biomed. Eng. 60, 954–9 (2013).

4. Shang, J. et al. A Single-Port Robotic System for Transanal Microsurgery-Design and Validation. IEEE Robot. Autom. Lett. 2, 1510–1517 (2017).

5. Shaikh, S. N. Natural orifice translumenal surgery: Flexible platform review. World J. Gastrointest. Surg. 91, 456–459 (2010).

6. Ferry, B., Gervasoni, D. & Vogt, C. Stereotaxic Neurosurgery in Laboratory Rodent. Stereotaxic Neurosurgery in Laboratory Rodent (2014). doi:10.1007/978-2-8178-0472-9.

7. Athos, J. & Storm, D. R. High Precision Stereotaxic Surgery in Mice. in Current Protocols in Neuroscience (2001). doi:10.1002/0471142301.nsa04as14.

8. Fornari, R. V. et al. Rodent Stereotaxic Surgery and Animal Welfare Outcome Improvements for Behavioral Neuroscience. J. Vis. Exp. (2012) doi:10.3791/3528.

9. Osten, P., Cetin, A., Komai, S., Eliava, M. & Seeburg, P. H. Stereotaxic gene delivery in the rodent brain. Nat. Protoc. (2007) doi:10.1038/nprot.2006.450.

10. Athos, J. & Storm, D. R. High Precision Stereotaxic Surgery in Mice. Curr. Protoc. Neurosci. 14, A.4A.1–A.4A.9 (2001).

11. Charles, P. & Watson, G. The rat brain in stereotaxic coordinates. (Elsevier, 2013).

12. Carter, M. & Shieh, J. Stereotaxic Surgeries and In Vivo Techniques. in Guide to Research Techniques in Neuroscience 73–88 (2015). doi:10.1016/b978-0-12-800511-8.00003-4.

13. Pak, N. et al. Closed-loop, ultraprecise, automated craniotomies. J. Neurophysiol. 113, 3943–3953 (2015).

14. Neurostar. https://neurostar.de/robot-stereotaxic/.

15. Brainsight. https://www.rogue-research.com/veterinary/microsurgical-robot/ Brainsight Vet Robot.

16. Snyder, W. E. & Qi, H. Fundamentals of computer vision. Fundamentals of Computer Vision (Cambridge University Press, 2017). doi:10.1017/9781316882641.

17. Ponce, J. & Forsyth, D. Computer vision: a modern approach. Computer (Pearson, 2012). doi:10.1016/j.cbi.2010.05.017.

18. Geng, J. Structured-light 3D surface imaging: a tutorial. Adv. Opt. Photonics 3, 128–160 (2011).

19. Hartley, R. I. & Sturm, P. Triangulation. Comput. Vis. Image Underst. (1997) doi:10.1006/cviu.1997.0547.

20. Stewart, D. A Platform with Six Degrees of Freedom. Proc. Inst. Mech. Eng. (1965) doi:10.1243/PIME_PROC_1965_180_029_02.

21. Dasgupta, B. & Mruthyunjaya, T. S. Stewart platform manipulator: A review. Mech. Mach. Theory (2000) doi:10.1016/S0094-114X(99)00006-3.

22. Szufnarowski, F. Stewart platform with fixed rotary actuators: a low cost design study. Fac. Technol. Bielefeld Univ. Ger. (2013) doi:10.1016/S0094-114X(99)00006-3.

23. Nanua, P., Waldron, K. J. & Murthy, V. Direct Kinematic Solution of a Stewart Platform. IEEE Trans. Robot. Autom. (1990) doi:10.1109/70.59354.

24. Cui, Y., Schuon, S., Thrun, S., Stricker, D. & Theobalt, C. Algorithms for 3D shape scanning with a depth camera. IEEE Trans. Pattern Anal. Mach. Intell. (2013) doi:10.1109/TPAMI.2012.190.

25. Izadi, S. et al. Kinect Fusion: Real-time 3D Reconstruction and Interaction Using a Moving Depth Camera. Proc. 24th Annu. ACM Symp. User interface Softw. Technol. - UIST’11 (2011) doi:10.1145/2047196.2047270.

26. Du, P. et al. From CUDA to OpenCL: Towards a performance-portable solution for multi-platform GPU programming. Parallel Comput. (2012) doi:10.1016/j.parco.2011.10.002.

27. Albrecht, O., Dondzillo, A., Mayer, F., Thompson, J. A. & Klug, A. Inhibitory projections from the ventral nucleus of the trapezoid body to the medial nucleus of the trapezoid body in the mouse. Front. Neural Circuits 8, 83 (2014).

