## Supplementary information for "Robotic Stereotaxic System based on 3D skull reconstruction to improve surgical accuracy and speed"

### 1. 3D skull reconstruction based on structured illumination and geometrical triangulation

#### Structured illumination to determine the image pixel indexes observing the same skull surface location

The surface of a small rodent's skull is relatively featureless and lacking in visual contrast. This makes it difficult for many 3D image reconstruction techniques to reconstruct the surface profile with intrinsic surface contrasts. For this reason, structured illumination was used in this project to superimpose external visual contrasts onto the skull surface, allowing 3D profile reconstruction with high spatial accuracies. The basic idea of structured illumination is that through projecting a series of vertical and horizontal lines onto the skull surface, unique binary spatial codes can be superimposed onto the skull surface for positional identification. With the unique spatial codes, images with different projection lines can be combined to identify the same spatial location on the skull to determine the image pixel indexes for the two cameras.

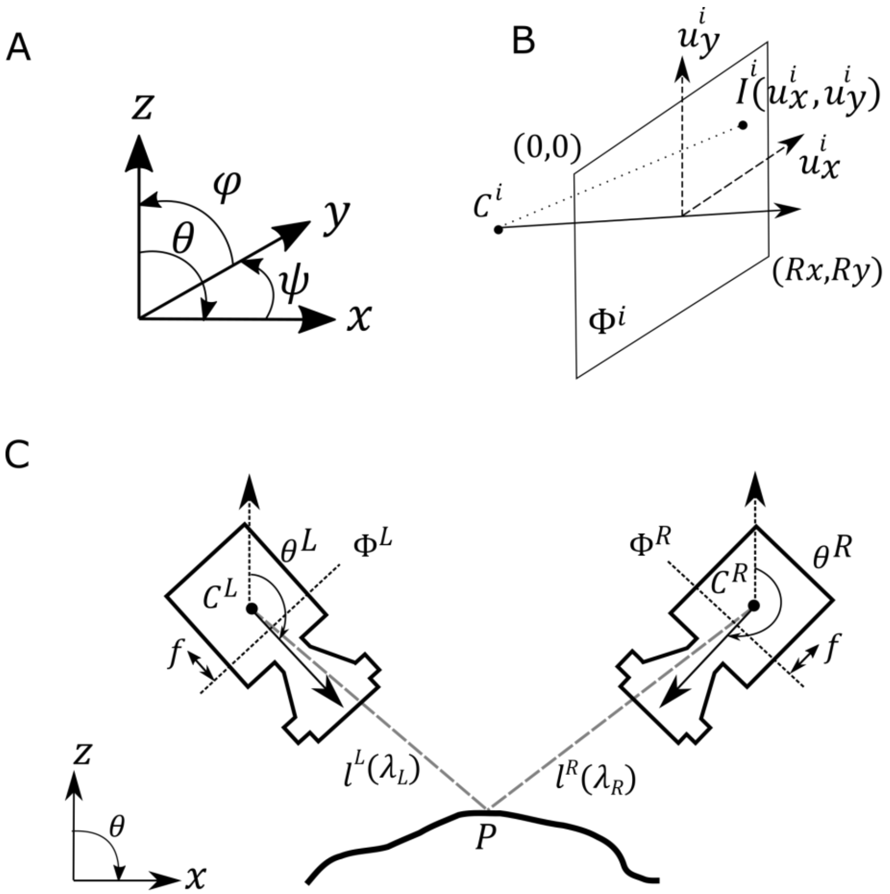


**Figure S1 A.** 3D profile reconstruction based on structured illumination and geometrical triangulation. **A.** Definition of the 6 degree-of freedom (6DOF). $\theta$ is the pitch angle rotating against the y-axis; $\psi$ is the yaw angle rotating against the z-axis; $\varphi$ is the roll angle rotating against the x-axis. **B.** A pixel with the image pixel index $(u_x^i, u_y^i)$ on the image plane $\Phi_\mathrm{i}$ of the 2D CCD camera with a camera center point $C^i$, where $i \in (L, R)$ to indicate the left or right camera. The origin of the image pixel index is at the upper left corner. **C.** The left and right CCD cameras lie on the x-z plane with a pitch angle $\theta^i$ focusing on the rodent's skull. Two lines $l^L(\lambda_L)$ and $l^R(\lambda_R)$ are projected from the two center points ($C^L$ and $C^R$) of the cameras going through the image planes to a cross point $P$ on the rodent's skull. The two cameras identify the image pixels corresponding to the same cross point $P$ using the binary spatial code constructed with structured illumination.

Using the determined image pixel indexes, the 3D coordinates of a skull surface location can be estimated using geometrical triangulation.

Forty black and white vertical and horizontal line patterns were projected onto the skull surface of a small rodent. In addition, one full-white and one full-black screens were also projected and imaged to allow monochromatic intensity superimposition onto the reconstructed skull profile for better visualization and removal of background image noise. Using this scheme of structured illumination, a maximum of $2^{40}$ unique binary codes can be generated to cover the entire skull surface of every location for positional identification.

**Geometrical triangulation to estimate 3D coordinates on the skull**

Figure S1A defines the roll ($\varphi$), pitch ($\theta$) and yaw ($\psi$) angles against the three translational axes $(x, y, z)$. Assume the center point of a CCD cameras is located at $C^i = \left(x_C^i, y_C^i, z_C^i\right)^T$ where $i \in (L, R)$ for the left or right camera. The camera has a calibrated image plane ($\Phi^i$) located in front of the center point $C^i$ with a distance equaling to the calibrated focal length $f^i$, as shown in Fig. S1C.

Images acquired by the cameras are formed by image intensities $I^i(u_x^i, u_y^i)$ and each pixel has an image pixel index $(u_x^i, u_y^i)$, as shown in Fig. S1B. Assume the left and right cameras are identical with the same image resolution of $(R_x, R_y)$, and the same pixel density of $(D_x, D_y)$ along the horizontal and vertical directions of the cameras, and the origin of the image pixel index is considered to be at the upper left corner, the actual physical displacement $dr^i = \left(dx^i, dy^i, dz^i\right)^T$ between the pixel $(u_x^i, u_y^i)$ and the center point of the camera center $C^i$ can be estimated by

$$dx^i = -D_x\left(\frac{R_x}{2} - u_x^i\right)$$

$$dy^i = D_y\left(\frac{R_y}{2} - u_y^i\right)$$

$$dz^i = f$$

The two cameras were setup to lie on the x-z plane of the laboratory coordinate, such that $\psi^i = \varphi^i = 0$, but with two nonzero pitch rotation angles ($\theta^L$ and $\theta^R$) focusing onto the animal skull. Therefore, a pixel $(u_x^i, u_y^i)$ on the camera $i$ can be translated to the laboratory coordinate $P^i$ by

$$P^i = C^i + Rot(\theta^i) \cdot dr^i$$

where $Rot(\theta^i)$ is the rotational matrix of the pitch angle $\theta^i$

$$Rot(\theta^i) = \begin{bmatrix} cos\theta^i & 0 & sin\theta^i \\ 0 & 1 & 0 \\ -sin\theta^i & 0 & cos\theta^i \end{bmatrix}$$

The center point $C^i$ and the pixel $P^i$ form a vector $v^i$ pointing toward the cross point $P$ on the rodent skull, such that

$$v^i = P^i - C^i = Rot(\theta^i) \cdot dr^i$$

In addition, the mathematical expression for the line $l^i(\lambda^i)$ on the camera $i$ to reach the cross point $P$ on the rodent's skull projecting along the vector $v^i$ is

$$l^i(\lambda^i) = C^i + \lambda^i \cdot v^i$$

where $\lambda^i$ is a parametric variable ranging from $-\infty$ to $\infty$ to indicate all the points along the projected line.

If two points are selected from the two projected lines – one point per line, the vector formed by these two points $(l^L(\lambda^L) - l^R(\lambda^R))$ cannot be perpendicular to both pointing vectors $v^L$ and $v^R$ of the two lines at the same time, unless when the two points are the same point crossing the two projected lines. Therefore, the

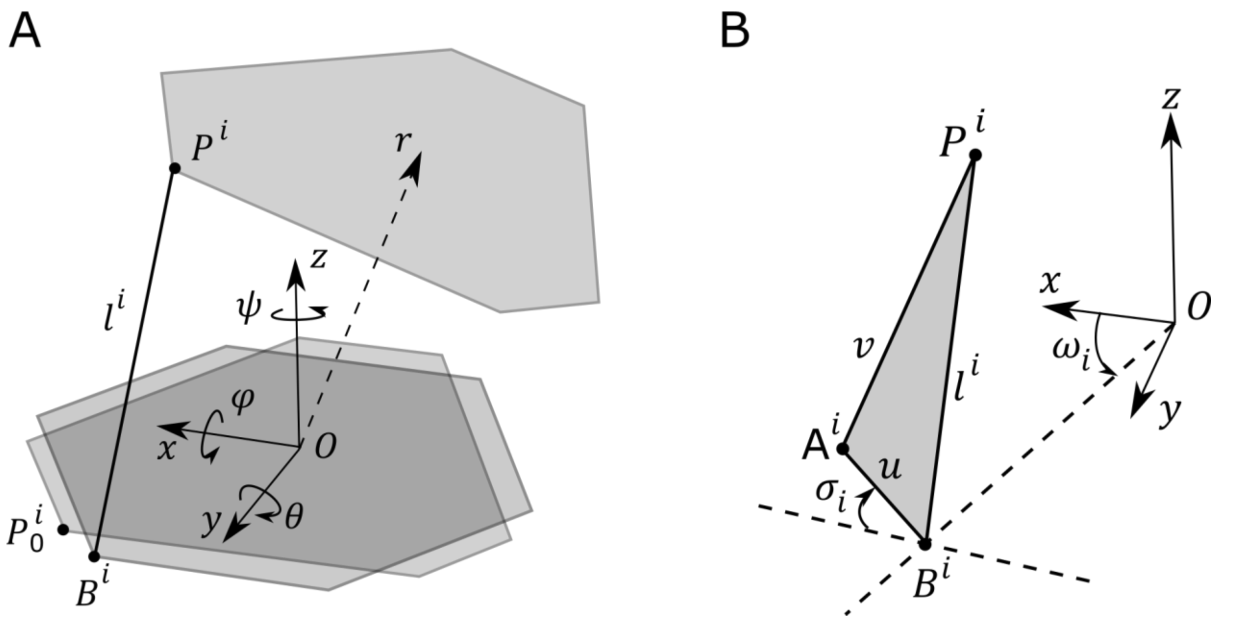


**Figure S2** Schematic drawing illustrating the top and bottom plates of the full 6DOF robotic platform for stereotaxic surgeries. A. The top and bottom platforms are connected by 6 arms and the arm length $l^i$ is determined by first rotating and then translating the top platform. B. The linear translation of the arm can be replaced by using a rotational servo with a rotational angle of $\sigma_i$.

cross point $P$ can be calculated by finding $\left(l^L(\lambda^L) - l^R(\lambda^R)\right) \cdot v^L = 0$ and $\left(l^L(\lambda^L) - l^R(\lambda^R)\right) \cdot v^R = 0$. Solving the two equations, the cross point $P$ can be estimated by

$$P = \frac{1}{2}(C^L + C^R + \lambda^L \cdot v^L + \lambda^R \cdot v^R)$$

where

$$\lambda^L = \frac{(v^L \cdot v^L)[(C^L - C^R) \cdot v^R] - (v^L \cdot v^R)[(C^L - C^R) \cdot v^L)]}{(v^L \cdot v^R)(v^L \cdot v^R) - (v^L \cdot v^L)(v^R \cdot v^R)}$$

$$\lambda^R = \frac{(v^L \cdot v^R)[(C^L - C^R) \cdot v^R] - (v^R \cdot v^R)[(C^L - C^R) \cdot v^L)]}{(v^L \cdot v^R)(v^L \cdot v^R) - (v^L \cdot v^L)(v^R \cdot v^R)}$$

### 2. Full 6DOF robotic platform based on Stewart design using digital servos

The top and bottom plates of the full 6DOF robotic platform can be designed to any shape or form as desired. However, in order to achieve full 6DOF positioning, installation of at least 6 pivots points are required for the top plate. Assuming one of the pivot points $i \in [1,6]$ has a spatial coordinate $P_0^i = \left(x_0^i, y_0^i, z_0^i\right)^T$ when the center points of the top and bottom plates are on top of each other, and supposing the center of the top plate were moved by a distance of $r = (x_r, y_r, z_r)^T$ and a rotation of $a = (\psi, \theta, \varphi)^T$ measured from the center of the bottom plate. The point $P_0^i$ will then be translated to $P^i = \left(x^i, y^i, z^i\right)^T$ with the following transformation equation

$$P^i = r + R \cdot P_0^i$$

where $R$ is the full rotational matrix such that

$$R = \begin{bmatrix} \cos\psi \, cos\,\theta & -\sin\psi\cos\varphi + \cos\psi\sin\theta\sin\varphi & \sin\psi\sin\varphi + \cos\psi\sin\theta\cos\varphi \\ \sin\psi\, cos\theta & \cos\psi\cos\varphi + \sin\psi\sin\theta\sin\varphi & -\cos\psi\sin\varphi + \sin\psi\sin\theta\cos\varphi \\ -\sin\theta & \cos\theta\sin\varphi & \cos\theta\cos\varphi \end{bmatrix}$$

Suppose the pivot point at the bottom plate has a spatial coordinate of $B^i = \left(x_B^i, y_B^i, z_B^i\right)^T$, the arm $i$ which connects the two pivot points $P^i$ and $B^i$ will have a length $l^i$ equaling to

$$l^i = \left|P^i - B^i\right|$$

Traditional Stereotaxic platform uses 6 high resolution linear actuators to extend or retract to the proper arm lengths $l^i$ to move the center of the top plate to a translational and rotational position $P_0^i = \left(x_0^i, y_0^i, z_0^i\right)^T$ and $r = (x_r, y_r, z_r)^T$. In this project, significant cost was saved by using rotational servo motors to replace the translational linear actuators to achieve the same arm lengths $l^i$ in which two arms with fixed lengths of $u$ and $v$ were connected by a pivot joint forming a triangle, as shown in Fig. S2B. In order to extend the hypotenuse arm of the triangle to have a length of $l^i$, a rotational angle $\sigma_i$ for the rotational servo motor is needed, and this rotational angle $\sigma_i$ can be calculated considering the 3D spatial position of the joint $A^i = \left(x_a^i, y_a^i, z_a^i\right)^T$ of the two short arms is

$$x_a^i = x_b^i + u\cos\sigma_i\sin\omega_i$$

$$y_a^i = y_b^i + u\cos\sigma_i\cos\omega_i$$

$$z_a^i = z_b^i + u\sin\sigma_i$$

where $B_i = \left(x_b^i, y_b^i, z_b^i\right)^T$ is the 3D spatial position of the rotational servo motor $i$ mounted on the base plate, and $\omega_i$ is the angle against the $x$ axis to the rotational servo motor $i$ on the $x - y$ plane of the base plate. The lengths of the three arms on the adjustable triangle can be calculated from the 3D spatial locations of $A^i$, $B^i$ and $P^i$.

$${l^i}^2 = \left(x_p^i - x_b^i\right)^2 + \left(y_p^i - y_b^i\right)^2 + \left(z_p^i - z_b^i\right)^2$$

$$u^2 = \left(x_a^i - x_b^i\right)^2 + \left(y_a^i - y_b^i\right)^2 + \left(z_a^i - z_b^i\right)^2$$

$$v^2 = \left(x_p^i - x_a^i\right)^2 + \left(y_p^i - y_a^i\right)^2 + \left(z_p^i - z_a^i\right)^2$$

The equations were combined and $x_a^i, y_a^i, z_a^i$ were substituted with their corresponding trigonometric relations, such that

$${l^i}^2 - v^2 + u^2 = 2u\left(z_p^i - z_b^i\right)\sin\sigma_i + 2u\left[\left(x_p^i - x_b^i\right)\sin\omega_i + \left(y_p^i - y_b^i\right)\cos\omega_i\right]\cos\sigma_i$$

Now define

$$\Lambda^{\mathrm{i}} = {l^i}^2 - v^2 + u^2$$

$$\Gamma^{\mathrm{i}} = 2u\left(z_p^i - z_b^i\right)$$

$$\Sigma^{\mathrm{i}} = 2u\left[\left(x_p^i - x_b^i\right)\sin\omega_i + \left(y_p^i - y_b^i\right)\cos\omega_i\right]$$

The angle for the digital servo $\sigma_i$ can therefore be determined by

$$\sigma_i = \sin^{-1}\frac{\Lambda^{\mathrm{i}}}{\sqrt{{\Gamma^{\mathrm{i}}}^2 + {\Sigma^{\mathrm{i}}}^2}} - \tan^{-1}\frac{\Sigma^{\mathrm{i}}}{\Gamma^{\mathrm{i}}}$$
